# Male-biased sexual selection persists across contrasting habitats in a dioecious plant

**DOI:** 10.64898/2026.02.19.706747

**Authors:** Camille Jolivel, Nina Joffard, Cécile Godé, Eric Schmitt, Isabelle De Cauwer

## Abstract

In most angiosperms, pollinators mediate access to sexual partners, making attractive floral traits key targets of sexual selection. Theory predicts stronger selection through the male function, due to greater dependence of male reproductive success on mate acquisition. However, habitat degradation may alter plant–pollinator interactions and disrupt sex-specific selection. In particular, reduced pollinator density may increase the dependency of female reproductive success on mate acquisition and strengthen selection on floral traits in both sexes. We tested this hypothesis in the dioecious species *Silene dioica* by measuring pollinator visitation, pollination service, and phenotypic selection on floral traits in both sexes across six populations located in either forest or anthropogenic habitats. Despite lower visitation in anthropogenic sites, pollination service was comparable between habitats with consistent selection on fertility-related traits in females. In contrast, we found stronger selection on a male trait likely promoting efficient pollen transfer in anthropogenic habitats, indicating a sex-specific response to land use change. Finally, male—but not female—reproductive success increased with the number of sexual partners in both habitats, supporting stronger sexual selection in males. More broadly, our results emphasize the need for a sex-specific perspective on floral trait evolution and demonstrate the persistence of male-biased sexual selection under environmental change.

## INTRODUCTION

A large majority of flowering plant species rely on insects for pollen transfer [1]. In these plants, reproductive success partly depends on complex interactions between plants and their pollinators, including the ability of plants to attract pollinators, and of pollinators to effectively remove, transport and deposit pollen onto receptive stigmas. Floral traits—such as flower number, floral colour, scent, morphology, or nectar production—can influence the frequency of pollinator visits [2–5] and/or the mechanical fit between flowers and pollinators [6–8], thereby affecting both pollen export (i.e., male function) and pollen receipt (i.e., female function).

Accordingly, floral traits are expected to be under pollinator-mediated selection, that is, selection arising from non-random pollinator visitation and handling that translates into differential reproductive success among plants. Pollinator-mediated selection has been widely documented across floral traits and species and is recognized as a major driver of angiosperm diversification [9–12]. Yet, most empirical studies focus on female reproductive success, typically quantified through individual seed production [9]. While this approach has yielded valuable insights into the evolutionary processes shaping floral traits, it provides an incomplete picture of the selection exerted on these traits, as it overlooks a substantial proportion of total reproductive output—either within hermaphroditic individuals or, in dioecious species, across the unmeasured sex.

If pollinator-mediated selection arises from variance in mate acquisition caused by differences in pollinator attraction among plants, then it can be regarded as a component of sexual selection [13,14]. Importantly, selection on floral traits is unlikely to be symmetric: it is predicted to differ between sexual functions both in strength and in the traits targeted. Whether and how such selection differs between sexual functions depends on the basic asymmetries of sexual reproduction. Such asymmetry stems from anisogamy, the fundamental difference in gamete size and number between sexual functions. Because male function typically involves the production of numerous small gametes, whereas female function relies on fewer, more resource-intensive gametes, male reproductive success is predicted to be more strongly constrained by mate acquisition, while female reproductive success is predicted to be more often limited by resource availability [15–17]. In plants, which generally produce far more pollen grains than ovules [18], this asymmetry should translate into siring success increasing with continued pollinator visitation and mate acquisition, whereas relatively few visits and sexual partners may suffice to maximize seed production. From this logic follows a clear prediction: selection through the male function should act predominantly on traits related to pollinator attraction (e.g., corolla size), whereas selection via the female function should primarily target traits related to fertility, that is, traits affecting the number and quality of gametes produced (e.g., flower number, ovule production per flower or resources provisioned to each ovule). This asymmetry should also be reflected by steeper Bateman gradients—the slopes of the regression of reproductive success on mating success, that provide a trait-independent measure of the fitness gain per additional mating and thus of the potential strength of sexual selection—in males than in females, as documented in numerous animal species [19] and in a growing, albeit still limited, number of plant species (e.g. in a moss, [20]; in a seaweed, [21]), including a handful of angiosperms (e.g., in a wind-pollinated species, [22]; and in insect-pollinated species, [23,24]).

The prediction that both selection on attractive floral traits and Bateman gradients should be stronger through the male than through the female function is valid only under optimal pollination conditions, when receptive stigmas receive enough pollen to fertilize all available ovules. However, pollen limitation—whereby pollen receipt limits seed production—is common in nature [16,25,26] and can be exacerbated by anthropogenic disturbances, such as habitat fragmentation, changes in floral resource availability, or increased pesticide and herbicide use, which reduce pollinator abundance and diversity [27–29]. Under such conditions, female reproductive success, just like male reproductive success, may become constrained by pollinator availability, which ultimately determines access to compatible mates. Consequently, both sexual functions (or sexes, in dioecious species) should benefit from additional pollinator visits, leading to stronger selection on attractive floral traits and steeper Bateman gradients in both sexes, reflecting stronger reproductive gains with increased access to mates. A few empirical studies have examined the relationship between pollinator density and selection strength and confirmed that selection on floral traits becomes stronger under pollen limitation [30–34]. However, such studies have almost exclusively focused on the female function (but see [35]), whereas a comprehensive understanding of floral trait evolution under pollinator decline requires considering selection exerted through both sexual functions.

In this study, we addressed this gap in knowledge by examining whether land use change alters plant–pollinator interactions and selection patterns in natural populations of *Silene dioica*, a dioecious species. Specifically, we quantified and compared pollinator visitation rates, pollination service, selection exerted on floral traits and Bateman gradients in both sexes between two contrasting habitats, by sampling three populations in minimally disturbed forest environments and three in intensively managed agricultural landscapes. We expected lower pollinator visitation rates and reduced pollination service in the agricultural context. Consequently, we predicted stronger selection on floral traits in both females—where pollinator-mediated selection may arise—and males—where it may intensify—in the anthropogenic habitat, as well an increase in the strength of sexual selection in both sexes, as indicated by steeper Bateman gradients in anthropogenic sites for both males and females.

## MATERIALS AND METHODS

### 1. STUDY SPECIES

*Silene dioica* (L) Clairv. (Caryophyllaceae) is a herbaceous, perennial and dioecious angiosperm widely distributed in central and northern Europe [36]. The species is described as predominantly forest-associated but can be found along a gradient from semi-natural habitats such as forest clearings or woodland edges to more anthropogenic environments like road verges and hedgerows, provided that conditions remain sufficiently moist and partially shaded [37]. It has a generalist pollination system and is primarily pollinated by bumblebees, honeybees, butterflies and hoverflies, the first two being the most frequent [23,36]. This species exhibits sexual dimorphism in several floral traits linked to pollinator attraction, such as flower size and number, with males displaying more conspicuous inflorescences [13,36,38]. In northern France, the species’ flowering period spans from late April to mid-July, peaking in late May.

### 2. STUDY POPULATIONS

Based on the land use in the area surrounding each population in a one kilometer radius (i.e., corresponding to the typical foraging range of bumblebees [39], one of the most common *S. dioica* pollinators), three populations from forest habitats (labelled “F”), containing at least 90% forest and semi-natural habitats, and three populations from anthropogenic habitats (labelled “A”), containing more than 40% agricultural and urbanized habitats, were selected (see electronic supplementary material – Habitat characterization). Study sites were selected to minimize spatial clustering of populations within each habitat type (**Figure 1**).

**Figure 1.**
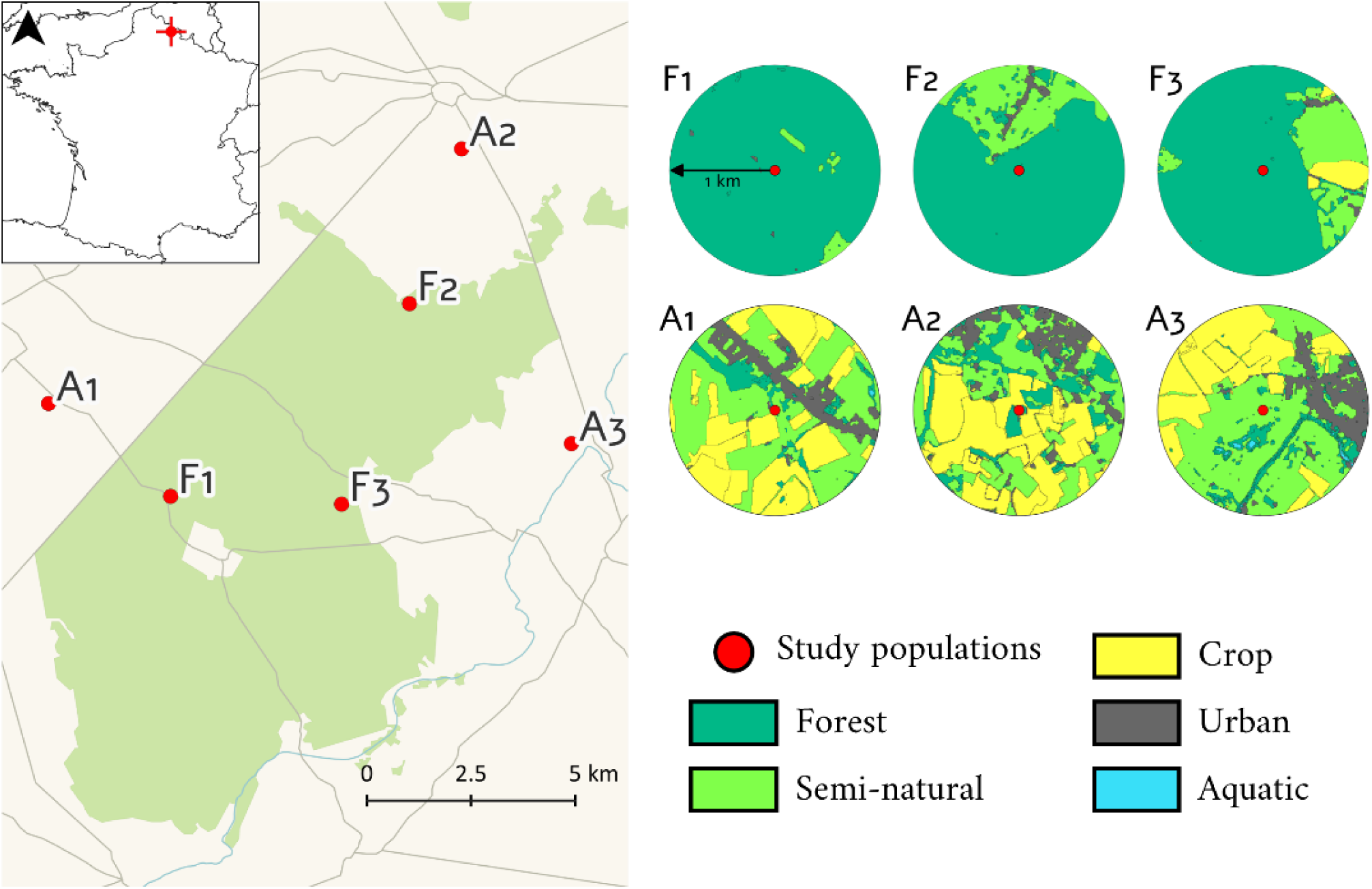
Location of the 6 populations (F: populations from forest habitats, A: populations from anthropogenic habitats) of *Silene dioica* sampled in and around the Mormal forest (Northern France) (left panel) and land use within a radius of 1 km around the center of each population (right panel).

In each population, 50 male and 50 female plants were randomly selected, ensuring individuals were at a comparable phenological stage (at or just before peak flowering) and aiming for a comparable density between the six populations (electronic supplementary material – Spatial configuration). These one hundred plants were individually tagged, and their geographical position was recorded (GPS 60 – Garmin). Outside of the tagged individuals, all *S. dioica* plants within the populations and within a 100-meter radius around the population had their inflorescences clipped before flowering, to ensure that the pollination events occurring during the study were primarily carried out by the fifty selected males.

In each of the six populations, flowering was monitored over ten days during the peak of the flowering period. On the first day, all pre-existing open flowers were marked with one colour of paint. During the experiment, newly opened flowers were marked with a different colour, allowing us to distinguish between flowers produced before and during the experiment. This procedure ensured that measurements of floral traits in males and females could be directly linked to individual reproductive success within the same time interval. The experiment was stopped after ten days because flower production became negligible, particularly in females which exhibit a shorter flowering duration than males [13].

### 3. FLORAL TRAIT MEASUREMENTS

On the first day of the experiment, for each individual, the number of open flowers was counted, and calyx height (mm) and corolla width (mm) were measured on two randomly selected flowers using calipers (precision: 0.01 mm). These two measurements were averaged before the analyses described below.

For female plants, the number of ovules per flower was estimated from three randomly selected fruits produced during the flowering survey, using an image analysis method developed by Barbot et al. [23]. This method involves dissecting the content of a fruit and digitizing it using a high-resolution scanner. Seeds and undeveloped ovules are differentiated based on their size and counted using an R script for image analysis (EBImage package in R, [40]). The number of ovules per flower corresponds to the sum of the seeds and undeveloped ovules, and was averaged over the three dissected fruits. For male plants, the number of pollen grains was estimated from two anthers collected from a nearly opened flower bud, using a particle counter (CASY Cell Counter & Analyzer, OLS) and following the protocol of Dufaÿ et al. [41]. This number was then multiplied by five to obtain the number of pollen grains per flower.

This set of floral traits was selected to include: (i) a trait whose role in pollinator attraction has already been established in *S. dioica* (corolla width, [42]); (ii) a trait putatively involved in pollinator fit (calyx height, which likely influences how deeply insects must enter the corolla to access nectar); and (iii) a trait directly related to fertility (number of gametes per flower). The last trait, number of open flowers, could affect individual reproductive success through its documented effect on pollinator attraction [2] and/or through its direct effect on gamete production [23].

### 4. POLLINATOR DIVERSITY AND DENSITY CHARACTERIZATION

To assess whether habitat type (F vs. A) affected pollinator density, pollinator observation sessions were conducted every other day during the duration of the experiment in each population. Each session lasted 20 minutes, always between 10:00 AM and 4:30 PM, and involved three independent observers, positioned at three distinct locations within the population. In total, 12 sessions were conducted in each population (three observers × four observation days). Each observer was responsible for a randomly chosen patch of focal plants (containing between 2 and 9 individuals, depending on their size and spatial arrangement) and recorded the identity of all pollinators visiting the patch (Orders: Coleoptera, Diptera, Hymenoptera, Lepidoptera, others), as well as the total number of flowers visited by each pollinator during its visit to the focal patch. A visit was defined as a contact between a pollinator and the reproductive structures of a flower. The total number of open flowers was recorded within each patch. Two estimators of pollinator density were derived from these observations: (i) the total number of visits and (ii) the flower visitation rate, calculated as the total number of visits divided by the total number of open flowers within the patch.

### 5. ESTIMATION OF POLLINATION SERVICE QUALITY

We performed a hand pollination experiment using plants kept in pots, in order to reveal potential differences in pollination service among populations independently of the quantity of resources available to the plants. Twenty-four female plants from a greenhouse collection were positioned within each study population and watered regularly during the experiment. These plants originated from seeds sampled in previous years in wild populations from the same area (in and around the Mormal State Forest) and grown in standardized conditions in a greenhouse at XXX University. Plants were randomly assigned to either an open pollination treatment (OP; 12 plants) or a hand pollination treatment (HP; 12 plants). Plants were arranged in groups of four along the length of the population, alternating between groups of four OP plants and groups of four HP plants (electronic supplementary material – Spatial configuration). Hand pollinations were carried out every other day, using pollen collected from fresh male flowers harvested from non-surveyed male plants located in neighbouring populations, to avoid removing flowers from the surveyed males. Hand pollinations involved rubbing the anthers of male flowers onto the stigmas of all open female flowers, ensuring that each stigma was fully saturated with pollen from at least two donors. At the end of the flowering survey, the 144 potted plants were returned to the greenhouse and monitored until fruit collection.

### 6. REPRODUCTIVE SUCCESS ESTIMATES

#### Females

For both sets of females (females from the six natural populations and potted), the number of fruits produced during the experiment was recorded. We also estimated the average number of seeds and ovules per fruit in three randomly selected fruits per female using the image analysis method described above. Then, for each female from natural populations, we estimated total seed production—a proxy of female reproductive success—by multiplying the average number of seeds per fruit by the total number of fruits produced. This estimation of female reproductive success was used for selection analyses. For each potted female, we calculated fruit set (number of fruits / number of flowers produced during the experiment) and seed set (number of seeds / number of ovules per fruit), as these indices allow us to assess whether pollen receipt was sufficient to maximize fruit initiation and seed production.

#### Males

Reproductive success of male plants was estimated using paternity analyses on a subsample of the seeds produced during the experiment. Total genomic DNA was extracted from dried leaf tissue for all adults (600 plants) and directly from seeds for 16 offspring sampled in eight randomly selected fruits per surveyed females (4,800 offspring in total, see [43] for seed DNA extractions) using the NucleoMag 96 Plant kit (Macherey Nagel, Oesingen, Switzerland) and an automated DNA purification station (KingFisher®, Thermo Fisher Scientific, Waltham, USA). PCR amplifications were performed on five nuclear microsatellites [23].

A spatially explicit mating model, developed by Oddou-Muratorio et al. [44] and modified by Tonnabel et al. [22] was used to estimate male reproductive success (*MRS*). This model uses information about the genotypes of the mothers, potential fathers and seeds, as well as the spatial distribution of individuals. This fractional paternity assignment method estimates the siring probability across all potential fathers for each genotyped seed. An average siring probability is calculated for all mother-potential father couples based on the 16 genotyped seeds. The number of offspring shared between a male and a female is then estimated as the product of the male’s siring probability and the female’s reproductive success (e.g., total number of seeds produced during the experiment, see previous section). The reproductive success of each male then corresponds to the sum of the seeds sired across all females in the population. Model parameters estimations are described in the electronic supplementary material (section Male reproductive success).

### 7. MATING SUCCESS ESTIMATES

Mating success (i.e., number of sexual partners) was estimated using the categorical paternity assignment method implemented in CERVUS 3.0.7 [45,46]. For each seed and each candidate male, a logarithm of odds (LOD) score was calculated, representing the likelihood ratio that the candidate is the true sire *versus* an unrelated male. To determine confidence in assignments, 10,000 simulations of reproduction events were performed to generate critical thresholds for delta LOD scores—the difference between the highest and second highest LOD scores—accounting for a 2% genotyping error rate and using an 80% confidence threshold. These thresholds were then applied to assign the most likely sire for each seed by comparing offspring genotypes to candidate males. Average assignment rate per population was 70% but varied among females. Therefore, to determine mating success a bootstrapping procedure was used: for each iteration, 8 seeds were randomly sampled for each female, a process that was repeated 100 times. The number of sexual partners for each male and female was then averaged across iterations.

### 8. STATISTICAL ANALYSES

#### Floral trait variation

Variation in floral traits between sexes and habitat types (F vs. A) was analysed using generalized linear mixed models (GLMMs), with a negative binomial distribution for the number of open flowers and a Gaussian distribution for calyx height and corolla width. Sex, habitat type and their interaction were treated as fixed effects. We also compared the number of ovules per flower and the number of pollen grains per flower between habitat types, using GLMMs with a Gaussian distribution, with habitat type treated as a fixed effect. Population was included as a random effect in all models.

#### Pollinator density

The total number of pollinator visits and the flower visitation rate were compared between habitat types (F vs. A) using a GLMM with a negative binomial distribution, including population and observation session as random effects.

#### Quality of pollination service

The quality of pollination service was compared between habitat types by assessing fruit set and seed set variation between OP and HP plants. To this end, we built GLMMs with a binomial distribution including pollination treatment (OP vs. HP), habitat type (F. vs A.) and their interaction as fixed effects, and population as a random effect. For the model focusing on seed set variation, female identity was also included as a random effect since seed set was measured on three fruits per plant. A significant interaction between the pollination treatment and habitat type interaction would indicate variation in the quality of pollination service between populations inhabiting forest and anthropogenic habitats.

#### Selection gradients

Within each population, selection gradients (*β*) were estimated following Lande & Arnold [47], using multiple regression analyses with relative reproductive success (individual reproductive success divided by mean reproductive success) as the response variable and standardized trait values (with a mean of 0 and a variance of 1) as explanatory variables (i.e., number of open flowers, number of gametes per flower, corolla width and calyx height). Selection gradients were estimated separately for males and females because reproductive success was estimated using different methods in the two sexes. Reproductive success was relativized and traits standardized separately for each population and sex.

To assess whether selection gradients varied between habitats, we used an ANCOVA including all traits, habitat type and interactions between each trait and habitat type as fixed effects and population as a random effect. A significant interaction between the effect of any trait and habitat type would indicate that the selection exerted on this trait differs between populations inhabiting forest and anthropogenic habitats.

Finally, because inbreeding depression can generate spurious correlations between traits and reproductive success when both are affected by inbreeding [48], we also estimated selection gradients including a proxy for individual inbreeding (internal relatedness [49]) as a covariate. This accounts for the fine-scale spatial genetic structure documented in *S. dioica*, likely resulting from limited gene flow and biparental inbreeding.

#### Relationship between traits, reproductive success and mating success

Structural equation modelling (SEM, [52]) was used to simultaneously quantify the direct effect of traits on reproductive success, the direct effect of traits on mating success (i.e., number of sexual partners), and the relationship between mating success and reproductive success, which here serves as a proxy for the Bateman gradient. If a trait influences reproductive success primarily through its effect on mating success, then this trait can be considered to be under sexual selection. One model was built for each population and sex, using the *piecewiseSEM* package in R. Reproductive success and mating success were relativized and traits standardized separately for each population and sex.

As a final step, we tested whether the relationship between reproductive success and mating success varied between habitat types using an ANCOVA with reproductive success as the response variable and mating success, habitat type and the interaction between the two as fixed effects. The number of open flowers and the number of gametes per flower were included as covariates to account for variation in fertility that could influence the relationship between reproductive and mating success, and population was included as a random effect. A significant interaction between mating success and habitat type would indicate that Bateman gradients differ between populations inhabiting forest and anthropogenic habitats. Analyses were conducted separately for males and females. Reproductive success and mating success were relativized, and traits were standardized separately for each population and sex.

All statistical analyses were performed using R (version 4.2.3). Normality of residuals and homogeneity of variance were verified for all linear models. Inspection of variance inflation factors (VIFs) in models estimating selection gradients indicated no problem of multicollinearity (all VIFs < 2, [53]). Overdispersion in generalized linear models with a negative binomial distribution was checked and found to be less than 2.

## RESULTS

### 1. FLORAL TRAIT VARIATION

The number of open flowers, corolla width and calyx height exhibited sexual dimorphism, with males having both more open flowers, and larger flowers (open flowers: *F*1,593 = 673.94, *p* < 0.001, corolla width: *F*_1,582_ = 228.07, *p < 0.001*, calyx height: *F*_1,582_ = 470.14, *p < 0.001,* electronic supplementary material – Floral trait variation). Additionally, the number of open flowers was significantly higher in the anthropogenic habitat compared to the forest habitat (*F*_1,593_ = 11.82, *p* < 0.001). The interaction between sex and habitat type was significant for calyx height only, with stronger sexual dimorphism in the anthropogenic habitat (*F*_1,582_ = 4.30, *p* < 0.05). The number of gametes per flower (i.e., number of ovules for females and number of pollen grains for males) did not differ between habitat types (Ovules: *F*_1,287_ = 2.51, *p* = 0.113, Pollen: *F*_1,294_ = 1.07, *p* = 0.301).

### 2. POLLINATOR DIVERSITY AND DENSITY

Across the whole dataset, the two most abundant pollinator orders were Hymenoptera (66% of all recorded visits), represented mainly by *Bombus* species, along with less frequent visitors such as honeybees and halictids, and Diptera (27% of visits), predominantly hoverflies (Syrphidae), with occasional occurrences of other families such as bee flies (Bombyliidae). Other observed orders included Coleoptera and Lepidoptera, but these were only recorded in populations F1 and A3, respectively. Apart from population A3, where hoverflies were the most frequent pollinators, the representation of the different pollinator groups appeared qualitatively similar across populations, with Hymenoptera accounting for the majority of visits, and no consistent differences observed between forest and anthropogenic sites (**Figure 2**).

**Figure 2.**
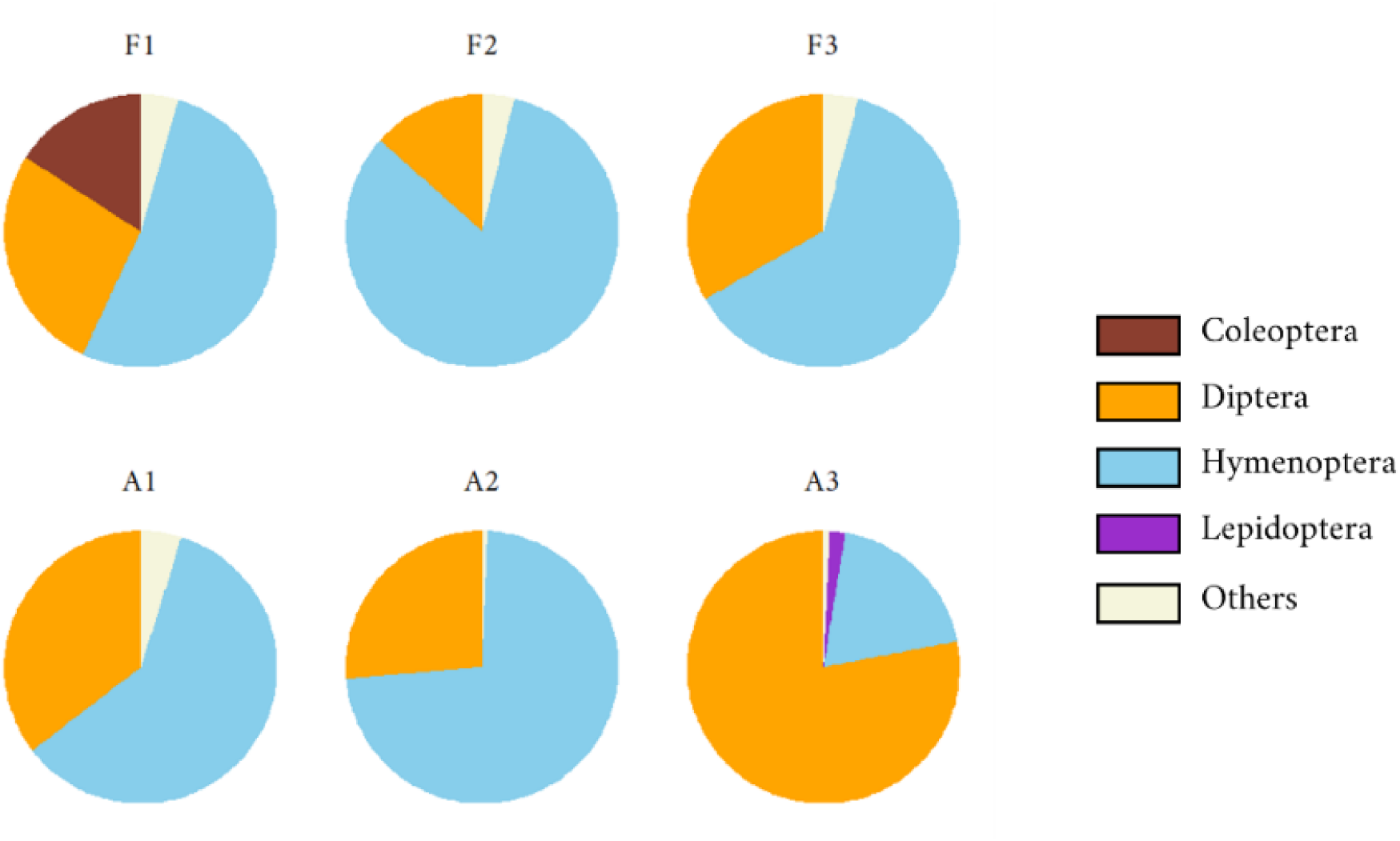
Relative frequencies of visits by different insect orders across the six natural populations (F: populations from forest habitats, A: populations from anthropogenic habitats).

The total number of visits per hour did not differ between populations located in forest and anthropogenic habitats (Forest: 51.00 ± 57.80 visits/hour, Anthropogenic: 75.25 ± 80.56 visits/hour, *F*_1,67_ = 0.50, *p* = 0.478). However, flower visitation rates (i.e., number of visits per flower per hour) were lower in populations located in anthropogenic habitats than in those located in the forest habitat (*F*_1,67_ = 3.99, *p* < 0.05), owing to the larger floral display of plants in the former. Flowers in the forest habitat received, on average, 1.4 times more visits than those in anthropogenic habitat (Forest: 0.660 ± 0.527 visits/flower/hour; Anthropogenic: 0.476 ± 0.594 visits/flower/hour).

### 3. QUALITY OF POLLINATION SERVICE

Fruit set was not affected by pollination treatment (*F*_1,135_ = 0.37, *p* = 0.544), habitat type (*F*_1,135_ = 1.36, *p* = 0.243), or their interaction (*F*_1,135_ = 1.06, *p* = 0.303) (electronic supplementary material – Pollination service quality). Similarly, seed set was unaffected by pollination treatment (*F*_1,391_ = 3.63, *p* = 0.060), habitat type (*F*_1,391_ = 0.29, *p* = 0.590), or their interaction (*F*_1,391_ = 0.28, *p* = 0.594; electronic supplementary material – Pollination service quality).

### 4. SELECTION GRADIENTS

#### Females

Selection gradients were significantly positive in all populations for the number of gametes per flower, and in all populations but one for the number of open flowers (F3, see **Figure 3**). Significant positive selection was also observed on calyx height in populations A1 and A3. As revealed by the significant interaction between trait and habitat type, selection on the number of open flowers differed between habitat types, being stronger in the anthropogenic habitat than in the forest one (γ_habitat x flowers_: 0.270 ± 0.089, *p* < 0.05). No effect of habitat was detected for the selection exerted on other traits (γ_habitat x gametes_: 0.006 ± 0.083, *p* = 0.942; γ_habitat x corolla_:-0.044 ± 0.087, *p* = 0.611; γ_habitat x calyx_: 0.138 ± 0.085, *p* = 0.104). Including individual internal relatedness in the model did not alter these conclusions (electronic supplementary material – Selection gradients with IR).

**Figure 3.**
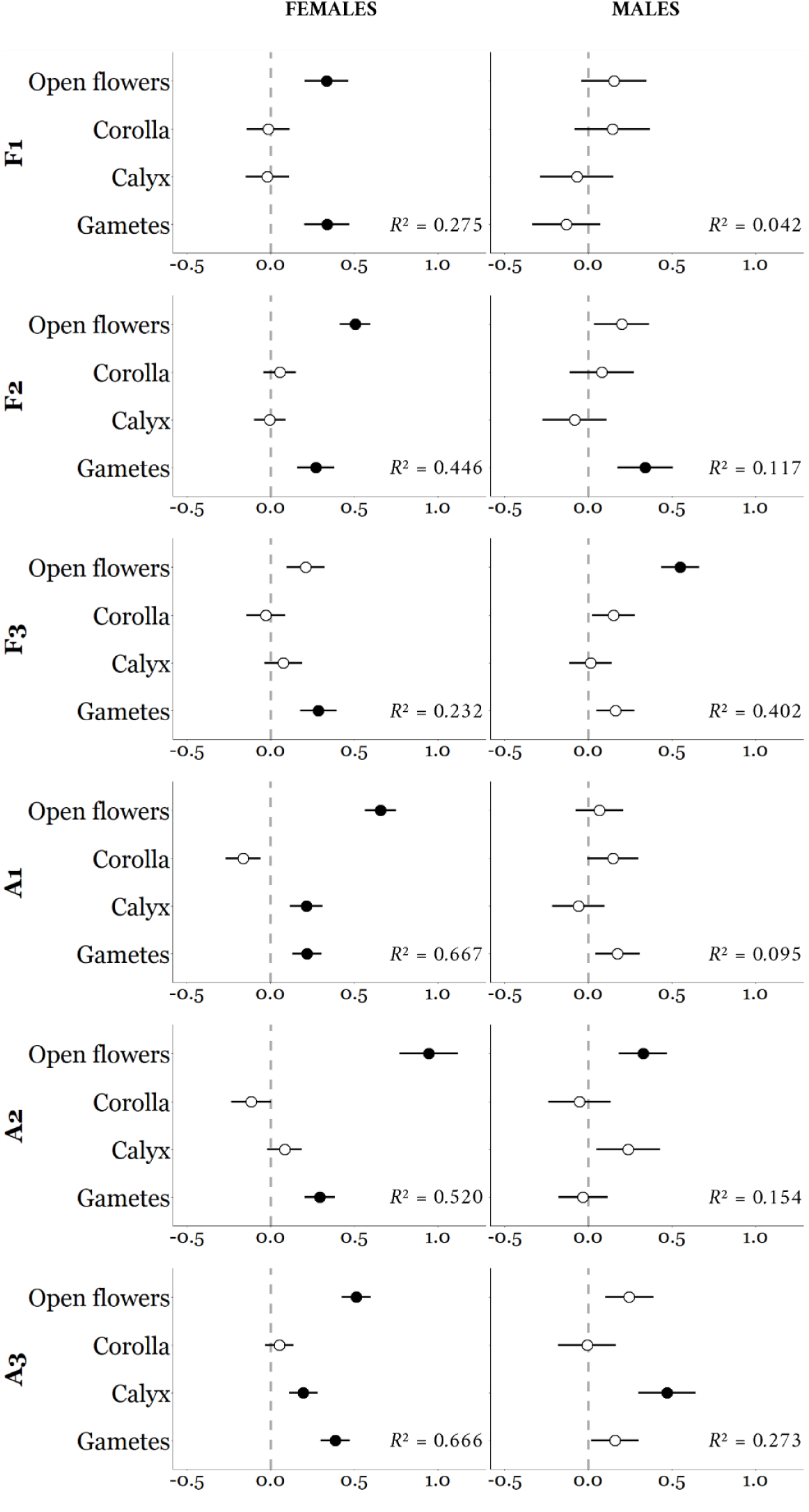
Selection gradients (± standard error) on floral traits for females (left panel) and males (right panel) in each natural population (F: populations from forest habitats; A: populations from anthropogenic habitats). Open flowers: number of open flowers; Corolla: corolla width; Calyx: calyx height; Gametes: number of gametes (ovules for females, pollen for males) per flower. Significant values are shown in black (*p* < 0.05).

#### Males

Positive selection was observed on the number of open flowers in populations F3 and A2, on the number of gametes per flower in population F2, and on calyx height in population A3 (**Figure 3**). Selection on calyx height differed between habitat types, with stronger selection in the anthropogenic habitat than in the forest one (γ_habitat x calyx_: 0.283 ± 0.141, *p* < 0.05). No effect of habitat was detected for the selection exerted on other traits (γ_habitat x flowers_:-0.032 ± 0.119, *p* = 0.787; γ_habitat x gametes_:-0.075 ± 0.121, *p* = 0.533; γ_habitat x corolla_:-0.112 ± 0.142, *p* = 0.427). Including internal relatedness in the analyses did not qualitatively change the patterns observed in selection gradients. However, internal relatedness had a significant negative effect on male reproductive success in one population (F2, electronic supplementary material – Selection gradients with IR).

### 5. RELATIONSHIP BETWEEN TRAITS, REPRODUCTIVE SUCCESS AND MATING SUCCESS

SEM allowed us to jointly estimate the effect of traits on reproductive and mating success, as well as the link between the two (i.e., proxy of the Bateman gradient), thereby disentangling direct and indirect effects of traits on reproductive success.

#### Females

Results obtained using SEM mirrored those obtained using the Lande & Arnold [47] method in all populations (**Figure 4**, electronic supplementary material – Structural equation modelling). A positive effect of the number of open flowers and the number of gametes per flower was detected in almost all populations, and a positive effect of calyx height was detected in populations A1 and A3. SEM analyses showed that these traits affected reproductive success in a direct way, not through an effect on mating success. Number of open flowers in population F1 and corolla width in populations F3 and A2 both had a positive effect on mating success. In these three cases, however, this positive effect on mating success did not translate into a positive effect on reproductive success, as there was no positive relationship between mating success and reproductive success. A negative relationship between the two was found in populations F1 and A1.

**Figure 4.**
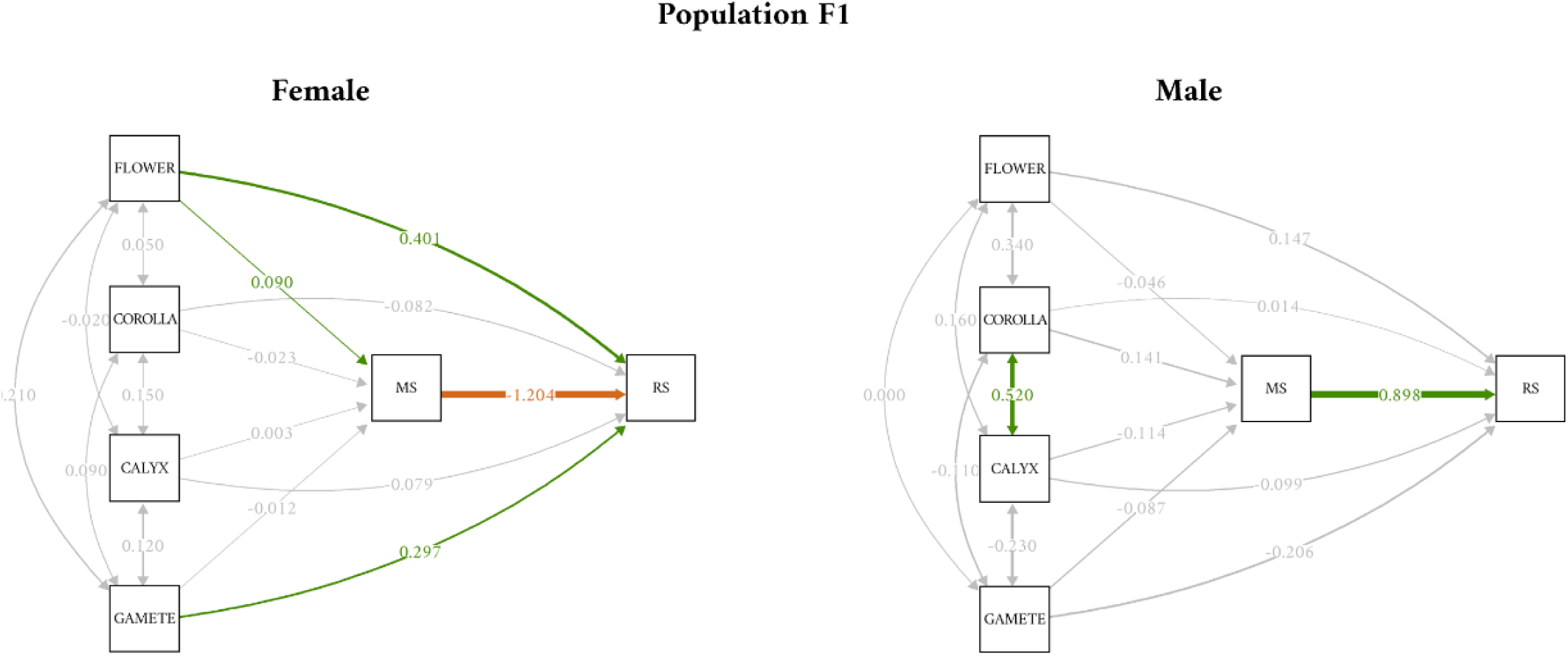
Example of path diagram of the structural equation model for population F1. Positives estimates are in green, negatives estimates are in orange, non-significant estimates in grey (*p* > 0.05). Line thickness represents the strength of the relationship between variables. Flower: number of open flowers; Corolla: corolla width; Calyx: calyx height; Gamete: number of ovules per flower; MS: mating success (i.e., number of sexual partners); RS: reproductive success (i.e., total seed production in females, total seeds sired in males).

#### Males

Overall, results obtained using SEM were largely congruent with those obtained using the Lande & Arnold [47] method (**Figure 4**, electronic supplementary material – Structural equation modelling). A positive, direct effect of number of open flowers and calyx height on reproductive success was detected in populations F3 and A3, respectively, while the previously observed positive effect of number of open flowers in population A2 appeared to be mainly indirect, as this trait had a positive effect on mating success, but no direct effect on reproductive success in this population. A higher number of flowers was also associated with a higher number of sexual partners in populations A3 and F3. In contrast to the pattern observed in females, in males, a positive relationship between mating success and reproductive success was detected in all but one population (F2), with no consistent difference in intensity between habitats.

The relationship between reproductive success and mating success was not affected by habitat, neither in males (*F*_1,268_ = 0.07, *p* = 0.789) nor in females (*F*_1,252_ = 0.12, *p* = 0.726) (electronic supplementary material – Habitat effects on Bateman gradients).

## DISCUSSION

In most angiosperms, access to sexual partners is mediated by pollinators, making floral traits key targets of selection. Sexual selection theory predicts stronger selection in males, yet environmental changes may alter this pattern. In particular, pollinator decline may increase intrasexual competition for mate acquisition and pollinator attraction, potentially leading to stronger selection on floral traits and steeper Bateman gradients in both sexes. In this study, we examined how habitat variation influences sex-specific selection on floral traits in the dioecious species *Silene dioica* by comparing populations located in forest and anthropogenic sites. Despite limited evidence for reduced pollinator activity and pollination service in anthropogenic sites, selection on a trait enhancing pollen transfer varied between habitats in males, whereas selection on female traits, primarily related to fertility, remained stable across sites. In line with sexual selection theory, male reproductive success showed a strong dependence on mating success, independently of habitat conditions, while no such dependence was observed in females.

### Habitat affected pollinator visitation rates but not pollination service

Pollinator visitation rates were significantly higher in the forest habitat, with flowers receiving, on average, 1.4 times more visits than flowers from anthropogenic sites, whereas total visit number did not differ between habitats. This suggests that pollinator density does not vary markedly between habitats, and that the reduced flower visitation rate observed in anthropogenic sites can instead be attributed to variation in floral display size rather than to differences in pollinator density *per se*. Indeed, plants in anthropogenic habitats had significantly larger floral displays, possibly due to greater light availability in more open environments or to fertilizer diffusion.

Given the observed differences in visitation rates, we expected reduced pollination service quality in anthropogenic habitats—reflected by a stronger difference in reproductive success between open-and hand-pollinated plants [25,54]. Contrary to expectations, we found no significant difference in the fruit set or seed set of open-and hand-pollinated plants in either habitat, suggesting that seed production in potted plants was not limited by pollen receipt. Since potted plants have access to fewer resources than wild individuals due to limited soil volume, we cannot extrapolate these results to naturally growing plants. Nonetheless, our findings indicate that if differences in pollination service quality exist between habitats, they are not detectable in plants with small floral displays and are therefore likely to be modest.

Despite the deliberate selection of sites that were strongly contrasted in terms of land use, we therefore found no marked differences in pollinator density or in the quality of pollination service between forest and anthropogenic populations. In light of what is known about the effects of agriculture on pollinator communities [27–29,55] and pollen limitation, this result is somewhat unexpected and suggests that common, generalist-pollinated plant species can be relatively insensitive to habitat differences of the magnitude considered here. The presence of heterogeneous landscape features such as meadows and agricultural hedgerows, as well as the relatively short distances to forested areas (A1: 2.2 km, A2: 3.2 km, A3: 1.8 km) in the three anthropogenic sites may contribute to maintaining high pollinator densities despite surrounding agricultural pressure.

### Selection on fertility-related traits in females

In line with our observation that pollination service appears adequate in all populations in both habitats, the traits most important for female reproductive success are those directly associated with fertility. In our analyses, selection acted primarily on the number of ovules per flower (in all populations) and on the number of open flowers (in all but one population), both of which directly determine the number of seeds a female can produce. These patterns were highly consistent across populations and align with previous findings in experimental populations of *S. dioica* experiencing no pollen limitation [13], further supporting the idea that, under sufficient pollen receipt, female reproductive success is largely determined by fertility traits.

Interestingly, we detected a difference in the strength of selection on flower number between habitats, whereby selection was significantly stronger in anthropogenic populations. Although flower number is known to be a major determinant of pollinator attraction in *S. dioica* [2,23] and other species [3,56,57], we argue that the reported difference in visitation rates between habitats is unlikely to explain such a pattern. First, as noted above, the quality of pollination service did not differ between habitats, suggesting that the reduced visitation rate was not sufficient to impact female reproductive success, at least in potted plants. Second, SEM analyses showed that the effect of flower number on reproductive success was predominantly direct, that is, not mediated by enhanced access to mates. Moreover, in cases where flower number apparently facilitated mate acquisition, we did not detect any positive association between mating success and reproductive success. We therefore argue that the stronger selection on flower number observed in the anthropogenic habitat is not due to reduced pollinator visitation in the latter, but to differences in trait distribution between habitats. Specifically, the number of open flowers exhibited higher variance in anthropogenic than in forest habitats (electronic supplementary material – Floral trait variation). Because standardized selection gradients are calculated on variance-standardized traits [47], enabling meaningful comparisons of selection strength across traits and studies, differences in trait variance between habitats can influence the estimated strength of selection even when the underlying fitness function remains unchanged.

Because females typically produce a limited number of gametes, even a relatively low number of pollinator visits may be sufficient to maximize female reproductive success in animal-pollinated plants. The absence of a positive relationship between mating success and reproductive success across the six populations sampled here supports that female reproduction does not benefit from an increased number of sexual partners, and therefore from enhanced attractiveness to pollinators in our study populations. Unexpectedly, we even detected a negative effect of mating success on reproductive success in two populations, implying that females producing more seeds also exhibited lower paternal diversity. Such a negative relationship may stem from variation in the intensity of post-pollination processes among females due to variation in resource availability. Specifically, females with better access to resources, because of their genotype or micro-environment, may both produce more seeds and exhibit greater potential for female choice [58,59]. Clearly, more studies are needed to disentangle the respective roles of pollinator visitation patterns and post-pollination processes in shaping the relationship between mating success and reproductive success.

### Mate acquisition is the strongest predictor of male reproductive success

In contrast to females, male reproductive success is expected to continue increasing at higher visitation rates, as the production of numerous gametes should make additional pollinator visits and pollen export to multiple females consistently advantageous. In this study, we indeed found a positive and significant relationship between mating success and reproductive success (i.e., positive Bateman gradient) in five populations out of six, indicating that siring success increases with mate acquisition, in line with theoretical expectations [15] and previous empirical findings [22–24]. Furthermore, this relationship was found to be unaffected by habitat, underscoring the high potential for sexual selection in males, and the persistence of such male-biased sexual selection in the face of land-use change.

Yet, the traits included in this study proved to be poor predictors of male mating success. Specifically, the only trait found to be under sexual selection in some populations was flower number, which was previously shown to enhance mate acquisition in *S. dioica* [23]. However, by manipulating this trait, Barbot et al. [23] showed that such an effect was not due to increased pollinator attraction but rather to altered pollinator movements and pollen export dynamics. Moreover, in our study no correlation was found between corolla width—a trait known to be involved in pollinator attraction in the focal species [42]—and male mating success across the six sampled populations. These results suggest that differential pollinator attraction may not be the primary source of variation in male mating success, and consequently not the main driver of variation in siring success in *S. dioica*. Instead, variation in siring success may depend on differential performances at subsequent stages of the male fitness pathway [60]. In line with this hypothesis, we detected positive selection on calyx height in one population, a trait promoting efficient pollen removal in other species [61]. Interestingly, selection on this trait differed between habitats, being stronger in the anthropogenic than in the forest one. Because the distribution of this trait was the same in the two habitat types, we suspect that such a difference truly reflects variation in the selective pressures experienced by plants in each type of sites. Specifically, deeper corollas may enhance the likelihood of contact between pollinators and the lower whorl stamens, which protrude only slightly from the flower. Calyx height could become more critical under low visitation, as observed in anthropogenic habitats, potentially leading to stronger selection on floral characteristics that improve pollen transfer [34].

Despite the inclusion of calyx height, the explanatory power of the male traits considered in this study remained limited, as reflected by the markedly lower *R²* values for selection models run on males compared to females. While such low R² values could partly reflect the fact that male success is inferred by genotyping a subsample of the seeds, while female reproductive success is assessed directly by counting all seeds, they may also indicate that we did not consider the male traits most important for shaping variation in siring success, particularly traits affecting pollen competition (e.g., pollen size, [62]). In *S. dioica*, *Bombus* sp.—which are the most common pollinators of *S. dioica* in the wild—can transport substantial amounts of pollen [43], and stigmas in natural populations often receive pollen loads that are significantly larger than needed to fertilize all ovules (pers. obs.), suggesting ample opportunities for post-pollination processes to occur [63,64]. In line with this hypothesis, including internal relatedness improved model fit in males—but not females—and revealed a significant effect of relatedness on siring success in one population, but no effect on the traits considered in this study. This suggests that inbreeding depression could affect other traits, perhaps associated with higher competitiveness at the post-pollination stage [65]. Overall, our results indicate a strong potential for sexual selection in males, yet this selection does not primarily target the attractive traits included in this study. Because pollination service was generally high across our study populations, including in anthropogenic sites, male–male competition may instead be expressed at later stages of the male fitness pathway.

## Acknowledgements

Many thanks to T. Desort, N. Roscigni, L. Meunier, A. Duputié, E. Houzé and N. Faure for their help during the field season. We are grateful to L. Debacker and S. Flourez for laboratory assistance. We would like to warmly thank colleagues and collaborators who provided advice, feedback, and discussions on data analyses: M. Dufaÿ, E. Barbot, J. Tonnabel, J. Shykoff and P. David. We would also like to thank the municipal authorities and owners of agricultural land for allowing us to work on their ground. This work has been performed using infrastructure and technical support of the “Plateforme Serre, cultures et terrains expérimentaux - Université de Lille” for the greenhouse/field facilities.

## Fundings

The authors thank the Région Hauts-de-France, the Ministère de l’Enseignement Supérieur et de la Recherche and the European Fund for Regional Economic Development for their financial support to the CPER ECRIN program, as well as the French Agence Nationale de la Recherche which financially supported our study through the ANR-JCJC-EvoPoD (ANR-21-CE02-0011).

## Author contributions

Conceptualization: CJ, NJ, IDC; Data Curation: CJ; Formal Analysis: CJ; Funding Acquisition: IDC; Investigation: CJ, NJ, CG, ES, IDC; Methodology: CJ, NJ, IDC; Project Administration: CJ, NJ, IDC; Resources: CG, ES; Supervision: NJ, IDC; Validation: NJ, IDC; Visualization: CJ; Writing—Original Draft: CJ; Writing—Review and Editing: CJ, NJ, IDC.

## Notes

### Competing Interest Statement

The authors have declared no competing interest.

