## supplemental file for "Male-biased sexual selection persists across contrasting habitats in a dioecious plant"

### Electronic Supplementary Material

#### 1. Habitat characterization

**Text S1**. Populations of *S. dioica* containing at least 100 plants were identified in the study region, in and around the Mormal State forest, a large forest area located in northern France (50°12'24.76"N, 3°44'14.84"E). The land use in the area surrounding each population (one-kilometer radius) was characterized and quantified using QGIS software (version 3.30.0) and the GroupStats plugin (version 2.2.7), using cartographic data processed by CNES for the THEIA Data and Services Center (www.theia.land.fr) from Copernicus satellite imagery. Based on these land-use data, we selected populations located in the most contrasting habitat matrices possible: three populations in predominantly forested landscapes and three populations in predominantly agricultural landscapes. Habitat types are detailed in Table S1 and were grouped from different subcategories to calculate the percentage of each habitat type around the six selected populations (Table S2).

**Table S1**. Habitat types found in the study area (in and around the Mormal State Forest) and correspondence with the database established by the THEIA Data and Services Center (French national inter-agency organization dedicated to facilitating the use of satellite and in-situ data for the observation and study of land surfaces).

| Habitat types | THEIA |
|---|---|
| Forest | Deciduous forest |
| | Coniferous forest |
| Semi-natural | Meadow |
| Crop | Straw cereals |
| | Sunflower |
| | Corn |
| | Tubers / Roots |
| Urban | Sparse urban area |
| | Road |
| Aquatic | Water |

**Table S2**. Frequency of each habitat type within a one kilometer radius around the center of each of the six studied populations. Populations with names starting with "F" are located in largely forested areas, whereas populations with names starting with "A" are situated in predominantly agricultural/anthropogenic landscapes.

| Population | Forest | Semi-natural | Crop | Urban | Aquatic |
|---|---|---|---|---|---|
| F1 | 0.976 | 0.023 | 0 | 0.002 | 0 |
| F2 | 0.799 | 0.184 | 0 | 0.016 | 0 |
| F3 | 0.772 | 0.175 | 0.041 | 0.012 | 0 |
| A1 | 0.072 | 0.443 | 0.372 | 0.113 | 0.001 |
| A2 | 0.151 | 0.311 | 0.412 | 0.126 | 0 |
| A3 | 0.086 | 0.474 | 0.312 | 0.122 | 0.005 |

#### 2. Spatial configuration

**Figure S1.** Spatial distribution of individuals within each population (F: populations from forest habitats, A: populations from anthropogenic habitats). In the figure, males from natural populations are shown in black and females in grey. White and yellow squares indicate the positions of potted individuals placed within each population to assess pollination service quality. Each white square corresponds to a group of four open-pollinated individuals, and each yellow square to a group of four hand-pollinated individuals.

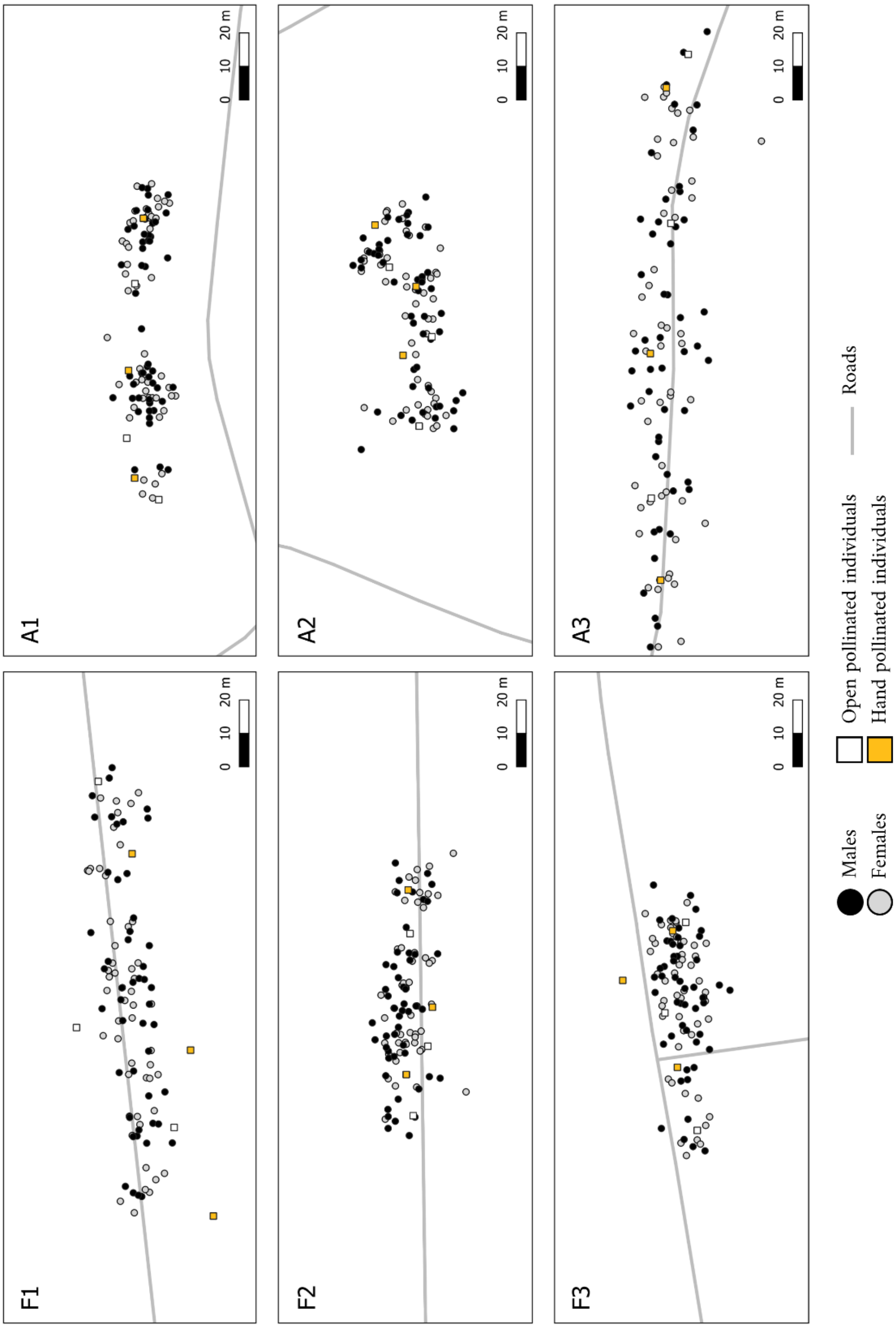

#### 3. Floral trait variation

**Table S3.** Trait mean ± SD for each sex and habitat. The table also shows the results of GLMs testing the effects of sex and habitat on flower number, corolla diameter, and calyx height, as well as the effect of habitat on gamete production (pollen and ovules), fruit set, seed set, and total seed number. Significant effects are indicated in bold. Flowers: number of open flowers; Corolla: average corolla width (mm); Calyx: average calyx height (mm); Ovules: number of ovules per flower; Pollen: number of pollen grains per flower; Fruit set: number of fruits divided by the number of flowers, Seed set: number of seeds divided by the number of ovules per fruit, average in three digitized fruits, Seeds: number of seeds produced.

| **Traits** | **Forest** | | **Anthropogenic** | | | | | | | |
|---|---|---|---|---|---|---|---|---|---|---|
| | **Females** | **Males** | **Females** | **Males** | $F_{sex}$ | $p_{sex}$ | $F_{hab.}$ | $p_{hab.}$ | $F_{sex*hab.}$ | $p_{sex*hab.}$ |
| **Flowers** | 2.63 ± 2.28 | 16.05 ± 15.93 | 6.04 ± 5.40 | 41.81 ± 47.04 | 673.94 | **< 0.001** | 11.82 | **< 0.001** | 1.28 | 0.257 |
| **Corolla** | 17.53 ± 2.94 | 20.61 ± 3.05 | 19.13 ± 3.19 | 22.78 ± 3.29 | 228.07 | **< 0.001** | 1.40 | 0.236 | 1.59 | 0.207 |
| **Calyx** | 12.78 ± 1.30 | 14.84 ± 1.17 | 13.08 ± 1.30 | 15.57 ± 1.71 | 470.14 | **< 0.001** | 0.83 | 0.361 | 4.30 | **< 0.05** |
| **Ovules** | 256.86 ± 65.96 | - | 275.94 ± 68.70 | - | - | - | 2.51 | 0.113 | - | - |
| **Pollen** | - | 70,426.06 ± | - | 87,162.95 ± | - | - | 1.07 | 0.301 | - | - |
| **Fruit set** | 0.966 ± 0.080 | - | 0.980 ± 0.056 | - | - | - | 0.70 | 0.404 | - | - |
| **Seed set** | 0.568 ± 0.198 | - | 0.600 ± 0.193 | - | - | - | 2.14 | 0.144 | - | - |
| **Seeds** | 2,376.29 ± 2,061.87 | - | 4,910.96 ± 4,921.96 | - | - | - | 3.63 | 0.057 | - | - |

**Table S4.** Trait standard deviation for each sex and population. Flowers: number of open flowers; Corolla: average corolla width (mm); Calyx: average calyx height (mm); Ovules: number of ovules per flower; Pollen: number of pollen grains per flower.

| | **Population** | **Flowers** | **Corolla** | **Calyx** | **Gametes** |
|---|---|---|---|---|---|
| Females | F1 | 1.71 | 2.56 | 1.26 | 61.14 |
| | F2 | 2.86 | 2.72 | 1.13 | 66.18 |
| | F3 | 2.02 | 2.88 | 1.11 | 68.43 |
| | A1 | 5.57 | 2.92 | 1.40 | 52.43 |
| | A2 | 5.79 | 2.35 | 1.27 | 65.95 |
| | A3 | 2.82 | 2.18 | 1.16 | 79.31 |
| Males | F1 | 8.78 | 2.30 | 1.26 | 22,543.14 |
| | F2 | 20.19 | 2.60 | 0.95 | 17,630.95 |
| | F3 | 14.59 | 3.04 | 1.20 | 12,887.58 |
| | A1 | 32.83 | 2.76 | 1.22 | 79,514.83 |
| | A2 | 63.60 | 2.74 | 1.08 | 19,210.36 |
| | A3 | 30.56 | 2.37 | 1.44 | 15,675.63 |

**Table S5.** Result for Levene tests for homogeneity of variance in females. Non-significant values after Bonferroni correction are indicated in grey ($p > 0.0055$). Blue boxes indicate that variance is greater in populations from anthropogenic habitats than in populations from forest habitats. Flowers: number of open flowers; Corolla: average corolla width (mm); Calyx: average calyx height (mm); Ovules: number of ovules per flower.

| **Females** | | | | | |
|---|---|---|---|---|---|
| **Flowers** | F2 | F3 | A1 | A2 | A3 |
| F1 | | | F1 < A1 | F1 < A2 | |
| F2 | | | | F2 < A2 | |
| F3 | | | F3 < A1 | F3 < A2 | |
| A1 | | | | | |
| A2 | | | | | A2 > A3 |
| **Corolla** | F2 | F3 | A1 | A2 | A3 |
| F1 | | | | | |
| F2 | | | | | |
| F3 | | | | | |
| A1 | | | | | |
| A2 | | | | | |
| **Calyx** | F2 | F3 | A1 | A2 | A3 |
| F1 | | | | | |
| F2 | | | | | |
| F3 | | | | | |
| A1 | | | | | |
| A2 | | | | | |
| **Ovules** | F2 | F3 | A1 | A2 | A3 |
| F1 | | | | | |
| F2 | | | | | |
| F3 | | | | | |
| A1 | | | | | |
| A2 | | | | | |

**Table S6.** Result for Levene tests for homogeneity of variance in males. Non-significant values after Bonferroni correction are indicated in grey ($p > 0.0055$). Blue boxes indicate that variance is greater in populations from anthropogenic habitats than in populations from forest habitats. Flowers: number of open flowers; Corolla: average corolla width (mm); Calyx: average calyx height (mm); Pollen: number of pollen grains per flower.

| **Males** | | | | | |
|---|---|---|---|---|---|
| **Flowers** | F2 | F3 | A1 | A2 | A3 |
| F1 | | | F1 < A1 | F1 < A2 | F1 < A3 |
| F2 | | | | F2 < A2 | |
| F3 | | | | F3 < A2 | |
| A1 | | | | | |
| A2 | | | | | |
| **Corolla** | F2 | F3 | A1 | A2 | A3 |
| F1 | | | | | |
| F2 | | | | | |
| F3 | | | | | |
| A1 | | | | | |
| A2 | | | | | |
| **Calyx** | F2 | F3 | A1 | A2 | A3 |
| F1 | | | | | |
| F2 | | | | | |
| F3 | | | | | |
| A1 | | | | | |
| A2 | | | | | |
| **Gametes** | F2 | F3 | A1 | A2 | A3 |
| F1 | | | F1 < A1 | | |
| F2 | | | F2 < A1 | | |
| F3 | | | F3 < A1 | | |
| A1 | | | | A1 > A2 | A1 > A3 |
| A2 | | | | | |

**Table S7.** Pearson correlation coefficients among floral traits, and between floral traits and internal relatedness, for males (upper triangle) and females (lower triangle) in each population (F: populations from forest habitats, A: populations from anthropogenic habitats), described using Pearson's correlation coefficient. Significant values after Bonferroni correction are indicated by stars (*: $p < 0.05$, **: $p < 0.01$, ***: $p < 0.001$). Open flowers: number of open flowers; Corolla: corolla width; Calyx: calyx height; Gametes: number of gametes (ovules for females, pollen grains for males) per flower; IR: Internal Relatedness.

| | | Open flowers | Corolla | Calyx | Gametes | IR |
|---|---|---|---|---|---|---|
| F1 | Open flowers | | 0.34 | 0.16 | 0.00 | -0.23 |
| | Corolla | 0.05 | | **0.52*** ** | -0.11 | 0.00 |
| | Calyx | -0.02 | 0.15 | | -0.23 | -0.06 |
| | Gametes | 0.21 | 0.09 | 0.12 | | 0.08 |
| | IR | -0.08 | 0.06 | -0.19 | -0.34 | |
| F2 | Open flowers | | 0.05 | -0.03 | -0.07 | 0.08 |
| | Corolla | 0.04 | | **0.51*** ** | -0.02 | -0.05 |
| | Calyx | -0.10 | 0.37 | | -0.11 | 0.09 |
| | Gametes | -0.24 | 0.27 | 0.07 | | 0.15 |
| | IR | -0.04 | -0.08 | 0.11 | 0.18 | |
| F3 | Open flowers | | 0.10 | -0.08 | 0.04 | -0.09 |
| | Corolla | 0.18 | | **0.45** ** | 0.12 | 0.34 |
| | Calyx | 0.09 | 0.28 | | 0.06 | 0.32 |
| | Gametes | 0.16 | 0.17 | 0.07 | | -0.01 |
| | IR | 0.16 | 0.01 | 0.13 | -0.06 | |
| A1 | Open flowers | | 0.15 | -0.24 | **0.37*** | -0.09 |
| | Corolla | 0.34 | | **0.55*** ** | 0.11 | 0.06 |
| | Calyx | 0.30 | **0.48** ** | | 0.08 | 0.20 |
| | Gametes | 0.14 | 0.33 | 0.11 | | 0.07 |
| | IR | 0.12 | 0.15 | 0.29 | -0.13 | |
| A2 | Open flowers | | 0.02 | 0.09 | -0.10 | 0.01 |
| | Corolla | **0.50*** ** | | **0.64*** ** | 0.11 | -0.21 |
| | Calyx | 0.17 | **0.44** ** | | 0.22 | -0.21 |
| | Gametes | 0.04 | 0.15 | 0.22 | | -0.09 |
| | IR | 0.01 | -0.10 | 0.15 | -0.15 | |
| A3 | Open flowers | | -0.09 | 0.09 | -0.02 | -0.20 |
| | Corolla | 0.16 | | **0.55*** ** | 0.11 | 0.02 |
| | Calyx | 0.21 | 0.14 | | 0.08 | 0.09 |
| | Gametes | 0.14 | 0.11 | 0.23 | | -0.15 |
| | IR | 0.01 | 0.08 | -0.13 | 0.30 | |

#### 4. Male reproductive success

**Text S2.** From the six sampled populations, multilocus genotypes of georeferenced maternal plants and their progeny were used in a fractional paternity assignments method (Oddou-Muratorio et al., 2018 ; Tonnabel et al., 2019) allowing us to estimate male reproductive success (*MRS*). This method also allows the estimation of the proportion of pollen originating from outside the population (i.e., the migration rate, *m*), as well as the parameters of the population-level pollen dispersal kernel. The kernel, modelled using a negative exponential power function (see Klein et al., 2006), describes the likelihood of fertilization as a function of distance. Its parameters include *b* ($b>1$ corresponding to a thin-tailed kernel, $b<1$ to a fat-tailed kernel, $b=1$ to an exponential kernel), a shape parameter describing how sharply siring probability declines with distance, and $\delta$, the pollen dispersal distance for each male. Model parameters were estimated with three Markov chain Monte Carlo (MCMC) simulations of 500 000 steps and a burn-in of 100 000 steps each. *MRS* and $\delta$ are two latent random variables, following Gamma distributions, with means and variances of ($1$, $\sigma_{RS}$) and ($\mu_\delta$, $\sigma_\delta$), respectively. We used uniform prior distributions for the parameters $\sigma_{RS}$, $\mu_\delta$, $\sigma_\delta$, $b$ and $m$ within the intervals [0.01,100], [1,100], [0,100], [0.1,10] and [0,1], respectively. The convergence of the Markov chains was assessed using three criteria : (i) the Gelman–Rubin diagnostic (< 1), (ii) the Geweke diagnostic (−2 to 2), and (iii) an analysis of chain autocorrelation (near to 0).

**Table S8.** Spatially explicit mating model parameters estimates for six populations of red campion (*S. dioica*). $\sigma_{RS}$: male reproductive success variance, *b*: shape of the pollen dispersal kernel, *m*: migration rate, $\mu_\delta$: mean pollen dispersal distance (m), $\sigma_\delta$: variance of pollen dispersal distance. Populations with names starting with "F" are located in predominantly forested areas, while populations starting with "A" are found in mainly agricultural areas.

| **Population** | $\sigma_{RS}$ | ***b*** | ***m*** | $\mu_\delta$ | $\sigma_\delta$ |
|---|---|---|---|---|---|
| F1 | 2.16 | 0.92 | 0.53 | 6.02 | 10.38 |
| F2 | 1.30 | 4.64 | 0.40 | 4.95 | 2.56 |
| F3 | 1.41 | 2.64 | 0.37 | 5.50 | 3.28 |
| A1 | 0.96 | 7.63 | 0.35 | 2.31 | 4.39 |
| A2 | 1.55 | 8.20 | 0.41 | 2.19 | 4.64 |
| A3 | 1.64 | 0.84 | 0.48 | 25.43 | 2.06 |

#### 5. Pollination service quality

**Figure S3.** Boxplots of fruit set (panel A) and seed set (panel B) for open pollinated (white) and hand-pollinated (orange) plants in each habitat.

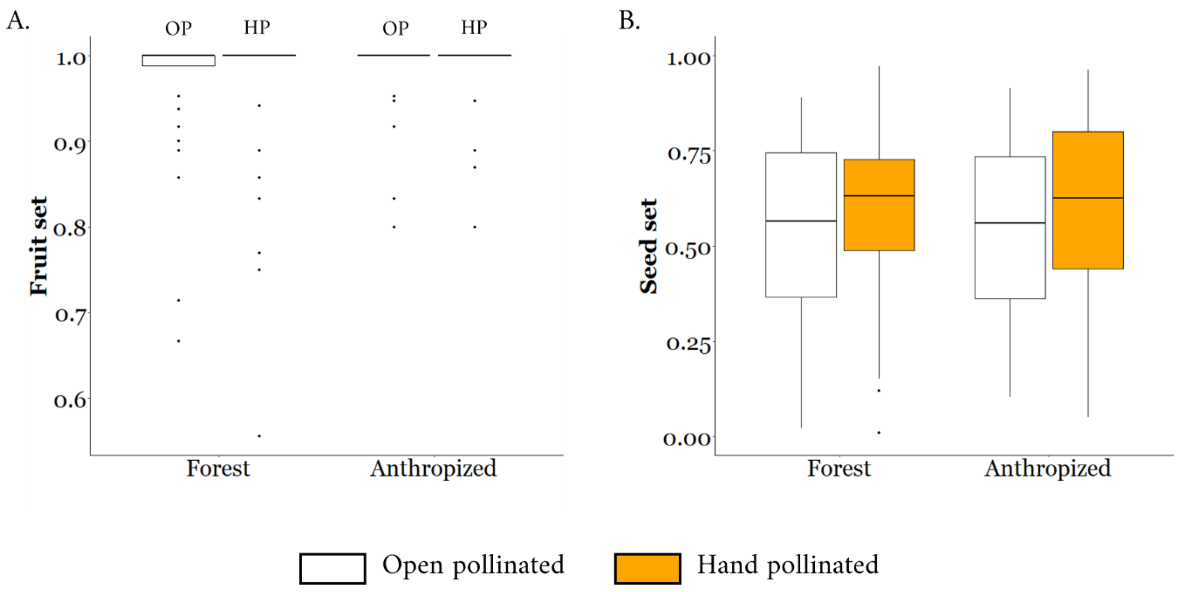

#### 6. Selection gradients with IR

**Figure S4.** Selection gradients (± standard error) on floral traits including internal relatedness (i.e., a proxy for individual inbreeding) as a covariate for females (left panel) and males (right panel) in each population (F: populations from forest habitats, A: populations from anthropogenic habitats). The slope of the relationship between reproductive success and internal relatedness is shown in the figure. Open flowers: number of open flowers; Corolla: corolla width; Calyx: calyx height; Gametes: number of gametes (ovules for females, pollen grains for males) per flower; IR: Internal Relatedness. Significant values are indicated in black ($p < 0.05$).

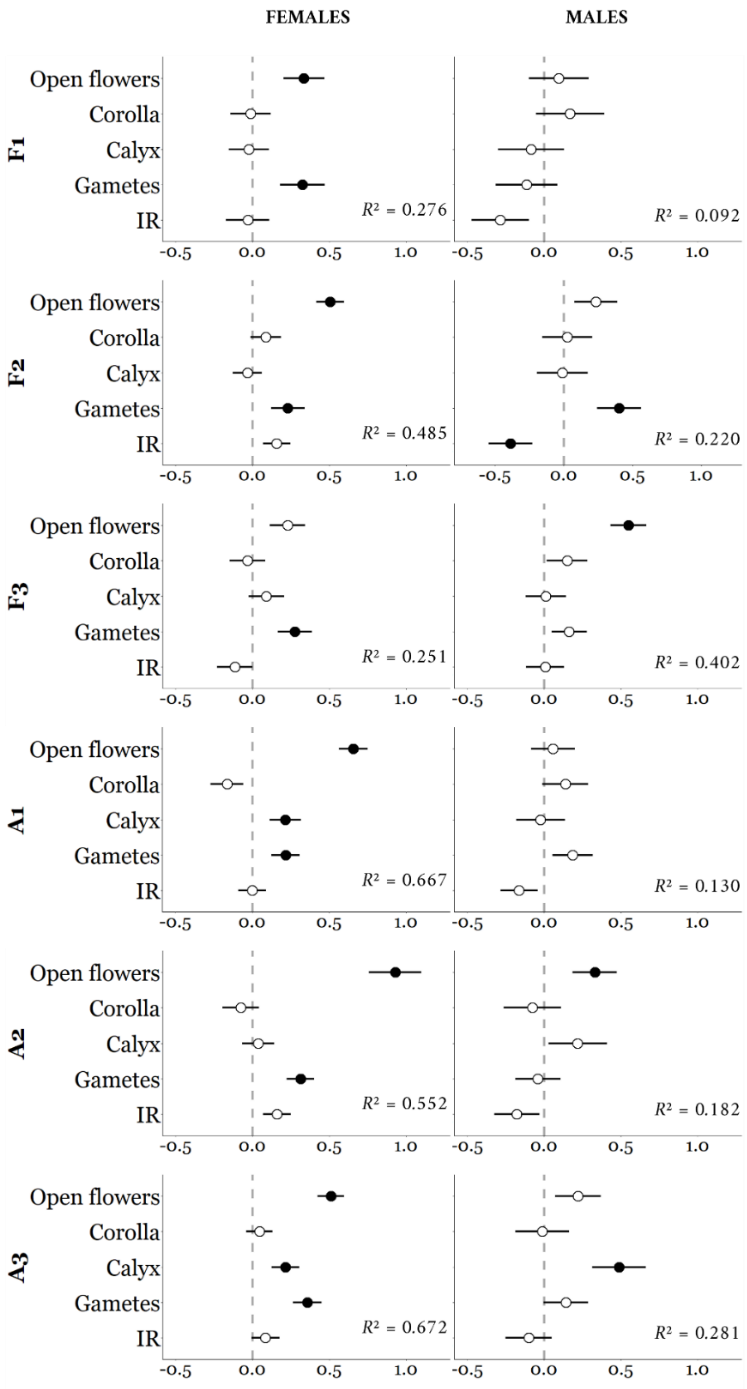

#### 7. Structural equation modelling

**Figure S5.** Path diagram of the structural equation model in females in each population (F: populations from forest habitats, A: populations from anthropogenic habitats). Positives estimates are in green, negatives estimates are in orange, non-significant estimates in grey ($p > 0.05$). Line thickness represents the strength of the relationship between variables. Flower: number of open flowers; Corolla: corolla width; Calyx: calyx height; Gamete: number of ovules per flower; MS: mating success (i.e., number of sexual partners); RS: reproductive success (i.e., total seed production).

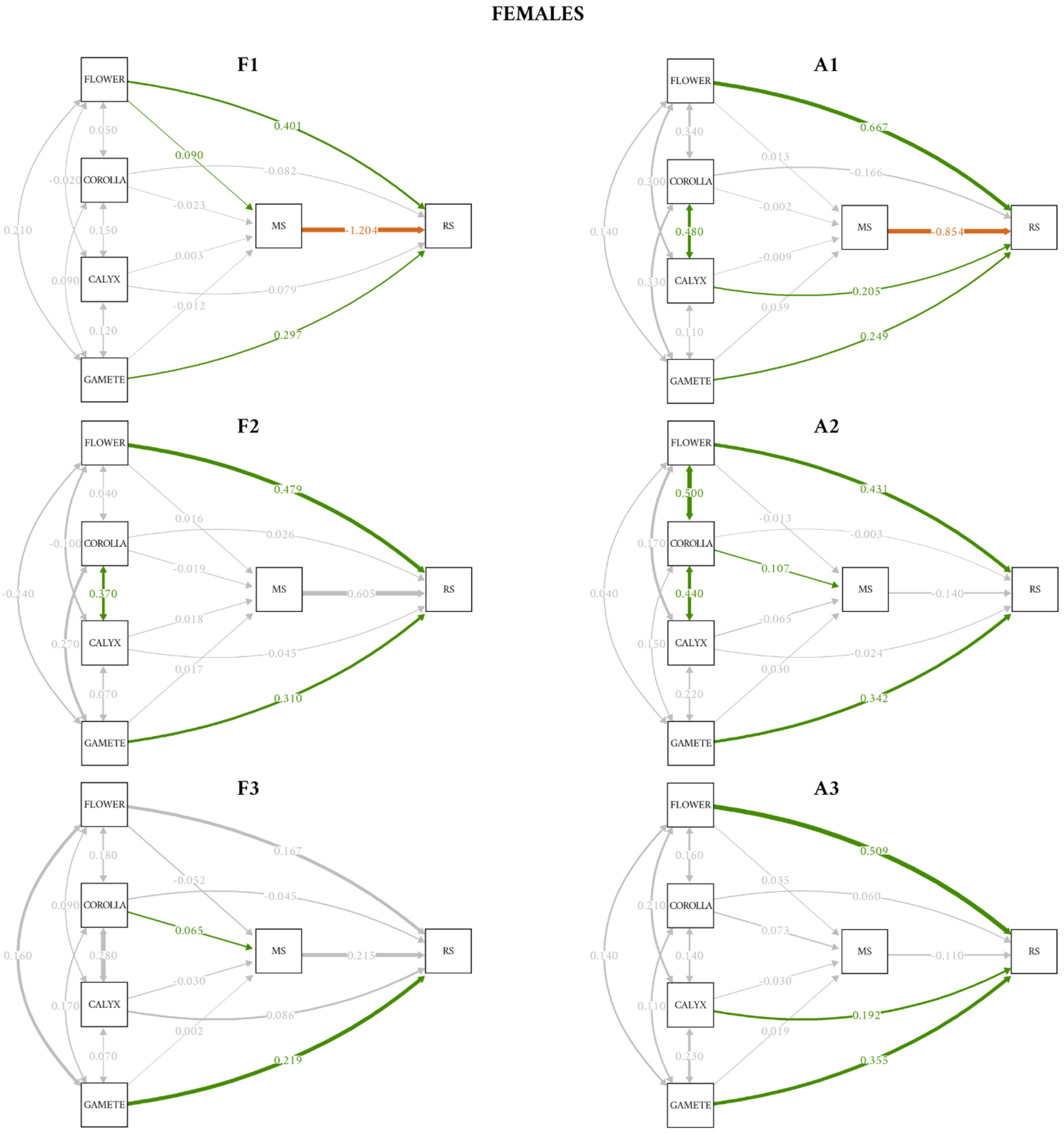

**Figure S6.** Path diagram of the structural equation model in males in each population (F: populations from forest habitats, A: populations from anthropogenic habitats). Positives estimates are in green, negatives estimates are in orange, non-significant estimates in grey (*p* > 0.05). Line thickness represents the strength of the relationship between variables. Flower: number of open flowers; Corolla: corolla width; Calyx: calyx height; Gamete: number of pollen grains per flower; MS: mating success (i.e., number of sexual partners); RS: reproductive success (i.e., total seeds sired).

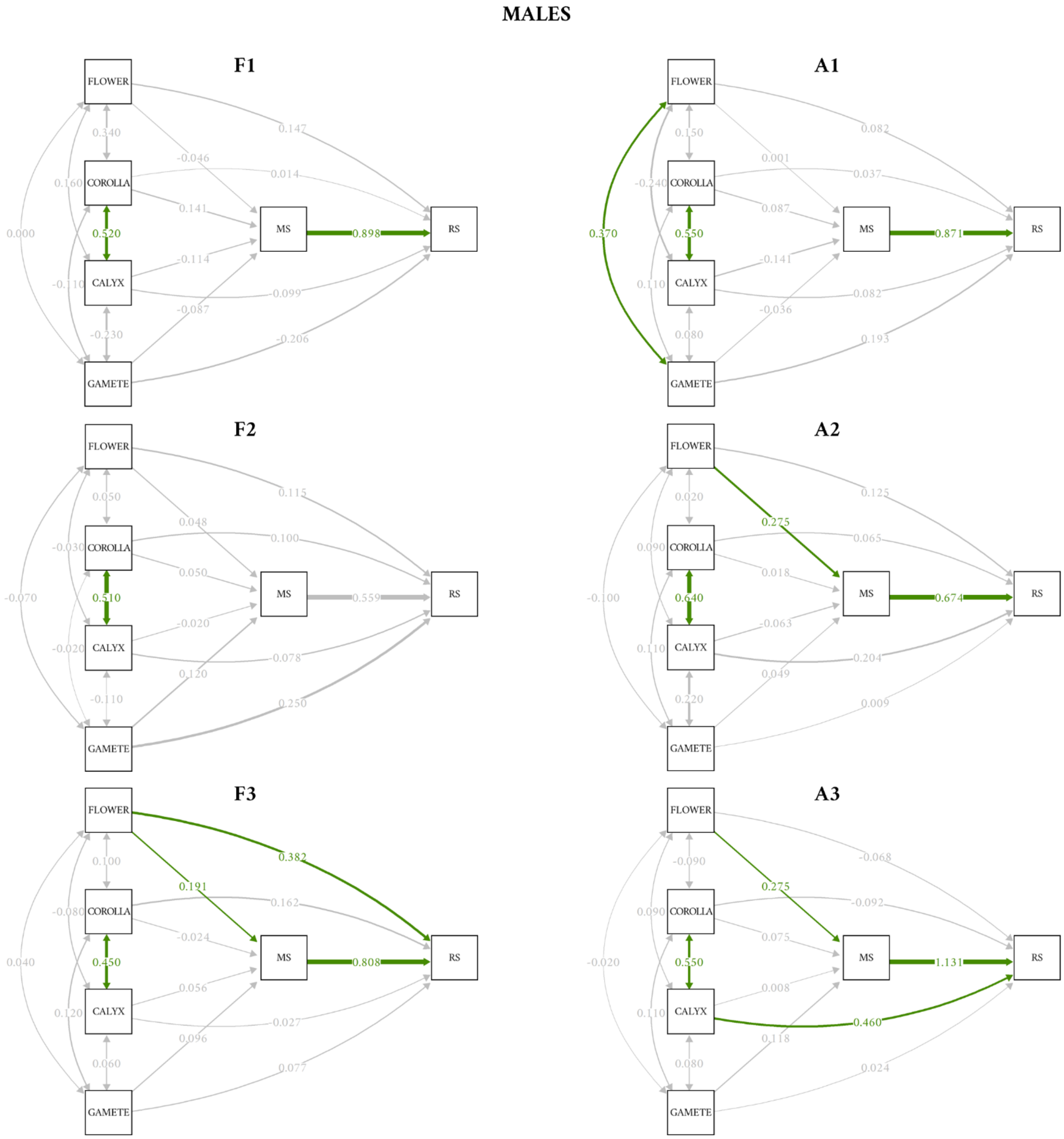

#### 8. Habitat effects on Bateman gradients

**Table S9.** Results of ANCOVA testing whether the relationship between reproductive success and mating success (MS) varies between habitat types. The number of open flowers (Flowers) and the number of gametes per flower (Gametes) were included as covariates, and population was included as a random effect. Forest habitat was used as the reference level in the model.

| | **Females** | | | **Males** | | |
|---|---|---|---|---|---|---|
| | Estimate | *F* | *p* | Estimate | *F* | *p* |
| MS | -0.245 | 0.20 | 0.288 | 0.813 | 85.31 | **< 0.001** |
| Habitat | -0.128 | 0.00 | 1.000 | -0.054 | 0.00 | 1.000 |
| Flowers | 0.434 | 108.84 | **< 0.001** | 0.156 | 7.79 | **< 0.01** |
| Gametes | 0.293 | 55.95 | **< 0.001** | 0.101 | 4.02 | 0.061 |
| MS*Habitat | 0.128 | 0.12 | 0.726 | 0.054 | 0.07 | 0.798 |
